# Diverging routes of oligodendrocyte recruitment in adaptive and regenerative myelination

**DOI:** 10.64898/2026.02.04.703508

**Authors:** Laura J Hoodless, Patricia Bispo, Elena MD Collins, Karl Baldacchino, Franziska Auer, Tim Czopka

**Affiliations:** Institute of Neuroscience and Cardiovascular Research, University of Edinburgh, United Kingdom; Simons Initiative for the Developing Brain, University of Edinburgh, United Kingdom; Institute of Neuronal Cell Biology, Technical University of Munich, Germany

## Abstract

Myelination by oligodendrocytes is dynamically regulated throughout life, supporting axon and circuit function during development, activity-dependent plasticity and repair after injury. How new oligodendrocytes are recruited from their lifelong pool of oligodendrocyte precursor cells (OPCs) in these distinct contexts remains unclear. Using high-resolution *in vivo* imaging in zebrafish, we directly compared activity-induced and demyelination-induced oligodendrogenesis under otherwise matched conditions. Although enhanced motor activity and demyelination both increased oligodendrocyte production, OPCs activated strikingly different transcriptional programmes. Demyelination, but not enhanced activity, induced acute expression of differentiation-associated genes, including the G-protein-coupled receptor Gpr17. Fate-tracking using a Gpr17 knock-in reporter revealed that all newly formed oligodendrocytes arise from Gpr17-positive states. In the healthy CNS, these states emerged exclusively through proliferation-linked OPC divisions, whereas demyelination unlocked a distinct, proliferation-independent conversion of homeostatic OPCs into Gpr17-positive, differentiation-primed cells. Our findings demonstrate the existence of diverging context-dependent routes of OPC recruitment engaged in adaptive and regenerative myelination.

## Introduction

Myelination by oligodendrocytes critically supports the health and function of central nervous system (CNS) axons through facilitating nerve conduction and provision of metabolites ^1^. It is now established that new myelin can dynamically form almost lifelong as part of ongoing and continued development ^2,3^, as well as in response to experience and neuronal activity in a process termed ’adaptive myelination’ ^4,5^. Likewise, new myelin can form in the regenerative process termed remyelination in disease and following injury ^6^. The myelin dynamics occurring in health and disease are largely driven by the differentiation of oligodendrocyte precursor cells (OPCs), which represent an abundant pool of specified, yet undifferentiated cells that reside throughout the grey and white matter lifelong ^7–9^. OPCs can differentiate to pre-myelinating oligodendrocytes which engage with surrounding target axons and, upon successful integration, mature to myelinating oligodendrocytes ^9–11^. While there is general consensus about the existence and necessity of these key stages, it remains largely unclear to what extent developmental, adaptive, and regenerative myelination share common or diverging mechanisms of oligodendrocyte recruitment from the pool of OPCs.

Work over the past years has revealed that OPCs are heterogenous, thus forming subpopulations with different properties and propensities to give rise to myelinating oligodendrocytes, depending on their state of lineage progression, intrinsic history and extrinsic local environment ^12–15^. This raises the possibility that, depending on the context, different OPCs may be recruited from their total pool to differentiate. Evidence exists that different pathways of oligodendrocyte may be employed in different contexts. For example, developmental and a range of remyelination studies support the concept that proliferative, ’activated’ OPCs strongly contribute to oligodendrocyte generation ^6^ . Likewise, direct fate tracking and live cell imaging studies in developing zebrafish and mice showed that new oligodendrocytes most frequently derive from recently divided OPCs ^14,16^.Other studies, however, have reported that new oligodendrocytes derive from the direct, proliferation-independent differentiation of OPCs, as seen for example in complex wheel running paradigms ^17,18^ as well as in sensory-driven myelination and remyelination in the adult mouse cortex ^19,20^. However, most existing studies either lack markers that allow for the discrimination of OPC subpopulations with different properties, or they lack the temporal resolution that is required to precisely pinpoint which OPCs give rise to new oligodendrocytes in a given context. Hence, whether these diverging observations reflect different mechanisms of recruitment from the OPC pool in healthy vs regenerative myelination, or other context-dependent variables such as age and CNS region studied (all of which likely affect the composition of the OPC pool), remains unknown.

Here, we tackle the question of how OPCs are recruited from their total pool in adaptive and regenerative developmental oligodendrogenesis, respectively. Taking advantage of the larval zebrafish model system, we carry out simultaneous *in vivo* imaging of multiple OPC markers in assays of adaptive and regenerative oligodendrocyte generation, thus presenting a model for highly-resolved analysis of oligodendrogenesis where only the stimulus to drive oligodendrocyte generation is altered. Using this model, we show that enhanced motor activity and demyelination both increase oligodendrocyte generation. However, transcriptome analysis revealed the differential induction of gene regulatory programmes in OPCs, where only demyelination led to an acute upregulation of markers indicative of differentiation. We generated a new knock-in reporter line for one of the markers, *gpr17*, which was selectively upregulated in demyelination and is actively discussed as an OPC subset marker in addition to labelling pre-myelinating oligodendrocytes ^21^.

Clonal timelapse analysis on the origin and fates of Gpr17-expressing oligodendrocyte lineage cells shows that it labels a subset of OPCs with a migratory phenotype, in addition to pre-myelinating oligodendrocytes. We further show that all newly differentiated oligodendrocytes emerge from a Gpr17-positive OPC state, and that the formation of new Gpr17-expressing OPCs occurred exclusively subsequent to OPC divisions in the healthy CNS. In contrast, demyelination unlocked a different mode of oligodendrocyte recruitment through direct, proliferation-independent acquisition of Gpr17-positive OPCs, thus fating these to differentiation. Therefore, our data reveal that adaptive and regenerative oligodendrogenesis occur via different routes of OPC recruitment.

## Results

### Enhanced motor activity as well as demyelination increase the rates of oligodendrocyte generation in the zebrafish spinal cord

To study context-dependent mechanisms of oligodendrocyte recruitment from the OPC pool, we developed independent assays that enhance oligodendrocyte differentiation in the zebrafish spinal cord in response to enhanced motor activity and demyelination, respectively (**Figs. 1, 2**). Two assays were used to enhance motor activity. In one assay referred to as the optical treadmill, enhanced zebrafish swimming was stimulated through the visual presentation of moving gratings (**Fig. 1a, Supplementary Movie 1**). These are perceived by the animal as whole field visual motion, which elicits an innate swimming behaviour called the optomotor response (OMR) ^22^. When these gratings were presented in a direction-changing manner, zebrafish continued to re-orient and follow the direction of flow, resulting in increased swimming behaviour (**Fig. 1a, Supplementary Movie 1)**. Concomitantly, using whole mount Hybridisation Chain Reaction (HCR) we detected increased expression of the immediate early gene *cfos,* which is commonly used as marker of enhanced neuronal activation (**Fig. 1b**). When we chronically enhanced motor activity on the optical treadmill between 4-5 days post fertilisation (dpf), we observed the increased generation of myelinating oligodendrocytes, confirming existing knowledge on activity-stimulated myelination (**Fig. 1c**). In addition to the optical treadmill, we also enhanced motor activity pharmacologically through the application of the voltage-gated potassium channel blocker 4-Aminopyridin (4AP). Analogous to the results from the optical treadmill and extending findings that we previously reported ^14^ , chronic, low-dose application of 4AP also enhanced swimming activity, *cfos* expression, and myelinating oligodendrocyte numbers (**Fig. 1d-f, Supplementary Movie 2).**

**Figure 1:**
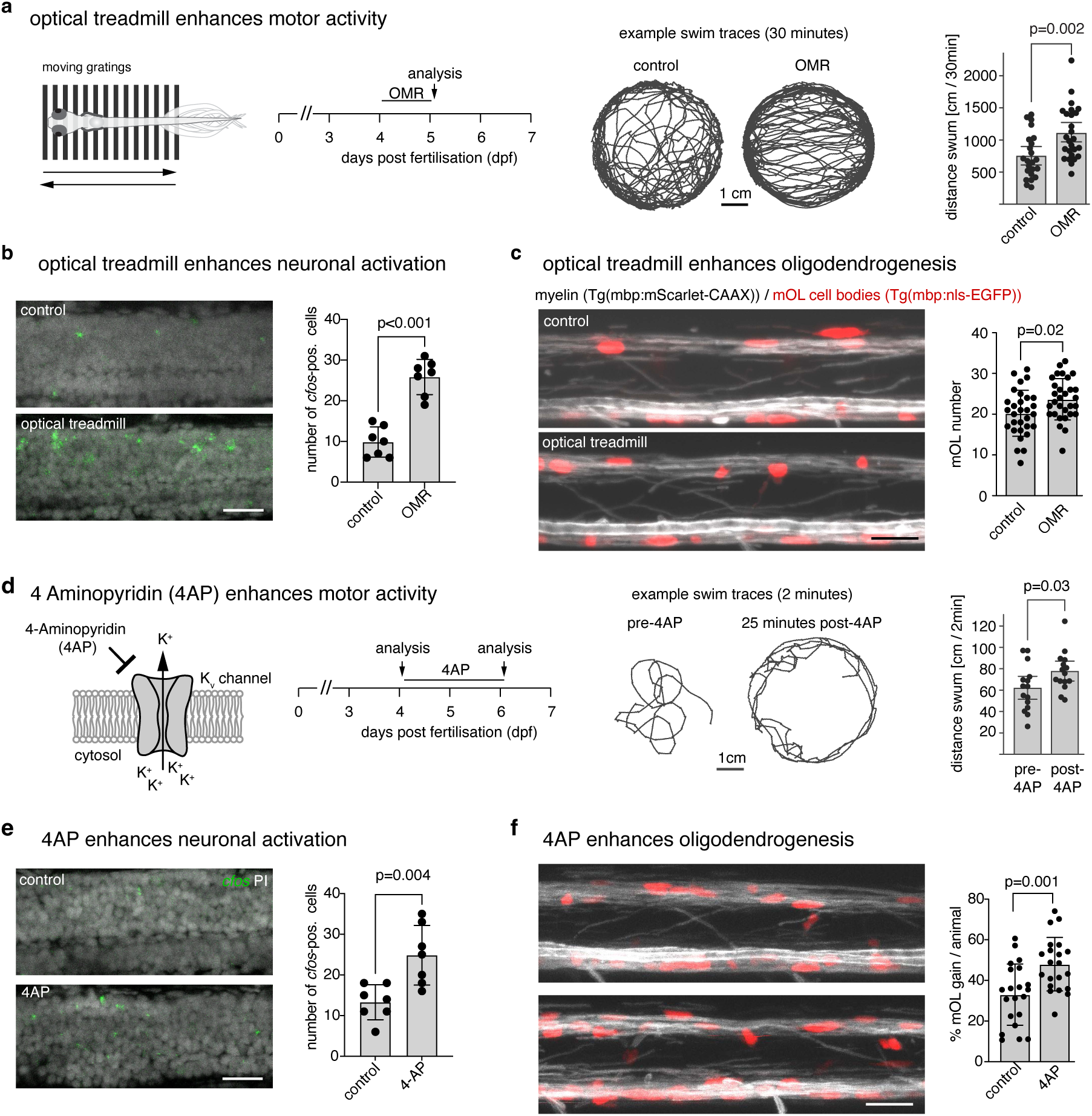
Assays to enhance oligodendrocyte generation through stimulation of motor activity. **a)** Schematic overview and timeline of the optical treadmill assay (left). To stimulate the optomotor response (OMR), fish were shown alternating moving gratings for 30-minute periods interspersed by 45-minute breaks. Right: Example traces of distance swum in control or OMR over 30 minutes. The distance swum was quantified in control and OMR between 5-7 d.p.f. Data are shown as mean ± 95% confidence interval (C.I.) cm swum in control *vs* OMR (743.5±146.4 *vs.* 1100.6±183.5; p=0.002, unpaired two tailed t-test, t=3.25, df=22); n=23/26 animals in control/OMR. **b)** Representative images of HCR *in situ* hybridisation for *cfos* in control and after the optical treadmill assay in lateral view of the zebrafish spinal cord. Graph shows quantification of the number of *cfos*-positive cells per field in the spinal cord. Data shown as mean ± standard deviation (S.D) cells/field in control *vs* OMR (9.9±3.6 *vs.* 25.9±4.21; p<0.001, unpaired two tailed t-test, t=7.412, df=12); n=7/7 animals in control/OMR. **c)** Representative images of Tg(mbp:mScarlet-CAAX), Tg(mbp:nls-EGFP) at 6 d.p.f. in control and after the optical treadmill assay (lateral view of spinal cord). Scale bar, 20 µm. Graph shows quantification of the number of myelinating oligodendrocytes (mOLs) per field in the dorsal spinal cord. Data are shown as mean ± S.D. cells/field in control *vs* OMR (20.2±5.6 *vs.* 23. 7±5.1; p=0.02, unpaired two-tailed t test, t=2.481, df=58); n=30/30 in control/OMR. **d)** Left: Schematic overview and timeline of the 4AP-mediated motor assay. Middle and right: Example traces of distance swum in control and 4AP over 2 minutes. Distances swum were quantified in the same animal pre-4AP and post-4AP at 5 d.p.f.. Data are shown as mean ± SD cm swum in control *vs* 4AP (61.7±21.5 *vs* 77.4±14.2 cm; p=0.03, paired two-tailed t test, t=2.49, df=14); n=15 animals assessed pre- and post-4AP in 1 experiment. **e)** Representative images of HCR in situ hybridisation for *cfos* in control and after 4AP treatment (lateral view of spinal cord). Graph shows quantification of the number of *cfos*-positive cells per field in the spinal cord. Data shown as mean ± SD cells/field in control *vs* 4AP (13.3±4.2 *vs.* 24.9±7.2; p=0.004, unpaired two-tailed t test, t=3.599, df=12); n=7/7. animals in control/4AP. **f)** Representative images of Tg(mbp:mScarlet-CAAX), Tg(mbp:nls-EGFP) at 6 d.p.f. in control and after 4AP treatment (lateral view of spinal cord). Graph shows quantification of the percentage of mOL gain per animal in the in the same region of the dorsal spinal cord between 4 and 6 d.p.f. Data are shown as mean ± S.D. cells/field in control *vs* 4AP (33.0±15.3 *vs.* 48.0±13.2; p=0.001, unpaired two-tailed t test, t=3.432, df=40. n=21/21 animals in control/4AP.

To contrast activity-induced oligodendrocyte recruitment in the healthy CNS with that occurring after demyelination, we adapted a chemogenetic ablation system where the transient receptor potential channel V1 (TrpV1) from *Rattus norvegicus* is selectively expressed in myelinating oligodendrocytes (**Fig. 2a**) ^23,24^. This ablation system is cell type specific because mammalian TrpV1 receptors but not endogenous zebrafish TrpV1 receptors are activated by capsaicin (Csn), leading to non-selective influx of cations only in the cells that express the mammalian receptor ^25,26^. Consequently, bath application of Csn to mbp:TrpV1-mScarlet transgenic animals results in the selective death of myelinating oligodendrocytes. In this assay, we observed the rapid recruitment of phagocytes within a few hours following induction of demyelination (**Fig. 2b**), and a near complete loss of myelinating oligodendrocytes within one day (**Fig. 2d, e).** Interestingly, in this model we also detected enhanced *cfos* expression levels one day post demyelination, suggesting that also this manipulation leads to some form of neuronal activation (**Fig. 2c**). Live cell imaging over 7 consecutive days revealed that macrophage numbers remained elevated until at least four days post demyelination, and returned to baseline levels by day 7 (**Fig. 2b**). At the same time, myelin returned to similar levels as pre-demyelination within four days (**Fig. 2d, e**). Developmental myelination is a protracted process in which new oligodendrocytes are added over long periods of time in most if not all species ^2,3,27^.

**Figure 2:**
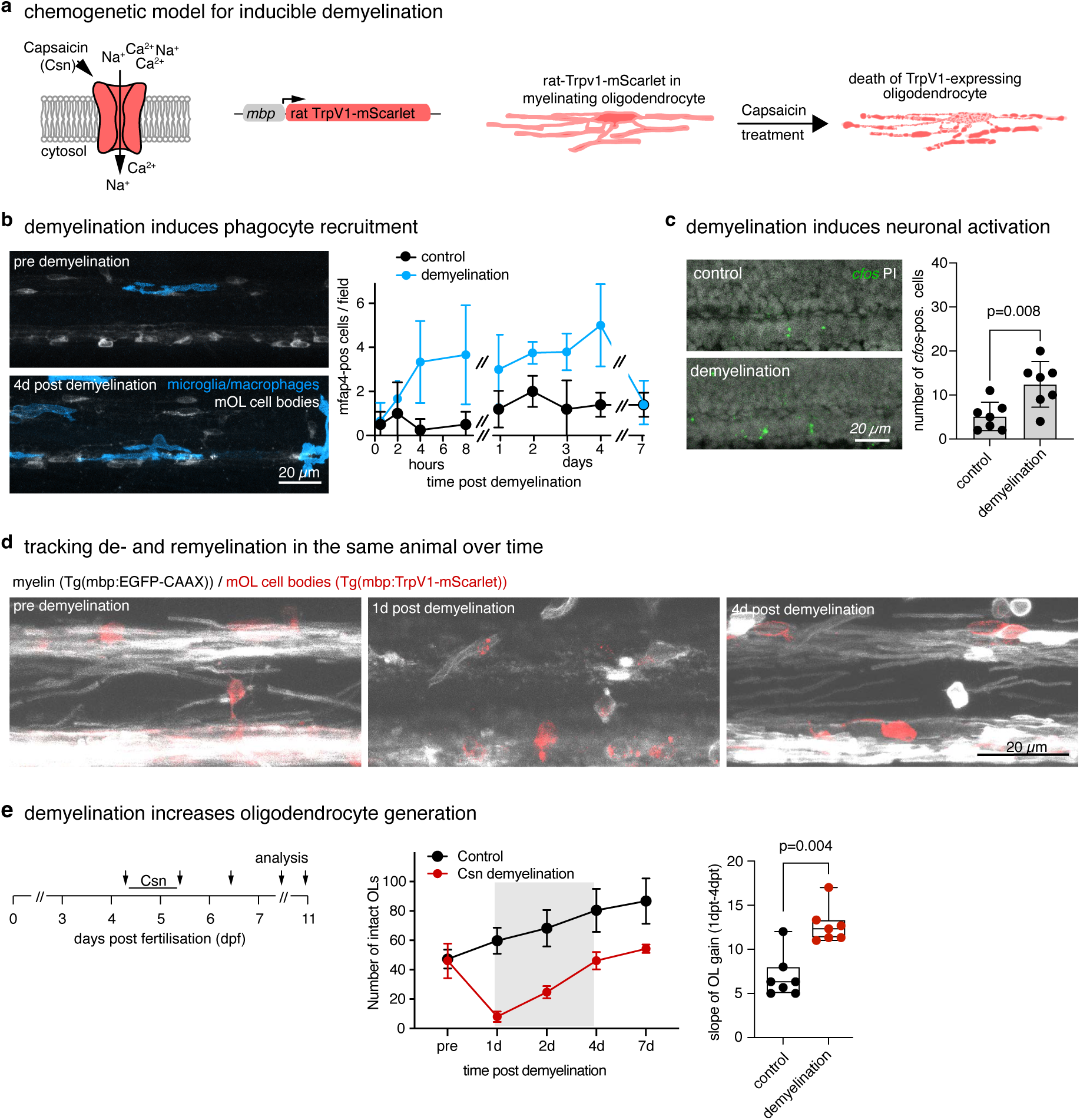
Enhancing oligodendrocyte generation through inducing demyelination. **a)** Schematic overview showing the principle of the mbp:TrpV1 demyelination assay. **b)** Representative images of Tg(mfap4:memCerulean), Tg(mbp:TRPV1-mScarlet) following demyelination (lateral view of spinal cord). Graph shows quantification of the number of mfap4+ microglia/macrophages at different timepoints after demyelination. Data shown as mean ± S.D cells/field in control *vs* demyelination (0.5h: 0.5±0.6 *vs* 0.7±0.8, 2h: 1.0±1.4 *vs* 1.6 ± 0.8, 4h: 0.3±0.5 *vs* 3.0±1.9, 6h: 0.5±0.6 *vs* 1.9±1.0, 8h: 0.5±0.6 *vs* 3.7±2.2, 1d: 1.2±0.8 *vs* 3.0±1.6, 2d: 2.0± 0.7 *vs* 3.8±0.5, 3d: 1.2±1.3 *vs* 3.8±0.8, 4d: 1.4±0.5 *vs* 5.0±1.9, 7d: 1.4±0.5 *vs* 1.5±1.0). n=5/7 animals in control/demyelination. **c)** Representative images of HCR in situ hybridisation for *cfos* in control and after demyelination (lateral view of spinal cord). Graph shows quantification of the number of *cfos*-positive cells per field in the spinal cord. Data shown as mean ± S.D cells/field in control *vs* demyelination (5.1 ± 3.2 *vs.* 12.4 ± 5.1, p=0.008, unpaired two-tailed t test, t=3.151, df=12); n=7/7 animals in control/demyelination. **d)** Example images taken of an individual Tg(mbp:TrpV1-mScarlet), Tg(mbp:EGFP-CAAX) transgenic animal before and at different time points after demyelination. Scale bar, 20µm. **e)** Left: Schematic showing the timeline of mbp:TrpV1-mediated demyelination assay. Middle: Graph depicting number of intact mOLs before and at various timepoints after demyelination. Data are expressed as mean ± S.D cells/field in control *vs* demyelination. (0h: 47.3±6.0 *vs.* 46.0±11.7, 1d: 59.7±8.7 *vs* 8.0±3.5, 2d: 68.3±11.9 *vs* 24.7±4.0, 4d: 80.4±14.5 *vs* 46.1±5.9, 7d: 86.7±15.2 *vs* 54.3±2.8). Right: Quantification of the slope of the line of mOL numbers between 1 d.p.t.-4 d.p.t. per fish, showing the rate of mOL gain. Data shown as median ± interquartile range (I.Q.R) in control vs demyelination (6.3±3.0 *vs* 12.3±2.0; p=0.004, Mann-Whitney U Test, *U*=3); n=7/7 animals in control/demyelination.

Consequently, we noted the continuous increase in myelinating oligodendrocyte numbers in control animals, and it should be noted that oligodendrocyte numbers in demyelination did not reach those seen in controls over the periods analysed (**Fig. 2e)**. However, when comparing the rate of oligodendrogenesis, we found an approximately two-fold increase over a 4-day period following demyelination compared to controls (**Fig. 2e)**. Therefore, this demyelination model, like the models of enhanced motor activity, enhances oligodendrocyte recruitment.

### RNA sequencing reveals activation of divergent transcriptional programmes in OPCs following demyelination and activity stimulation

To begin addressing whether OPCs are recruited by the same or by different mechanisms in response to demyelination and activity stimulation, we first sought to determine how their transcriptional programmes may change. Hence, we carried out bulk RNA sequencing of fluorescence-activated cell-sorted (FACS) OPCs from Tg(olig1:memEYFP) animals 1 day post demyelination and 4AP-mediated activity stimulation, respectively (**Fig. 3a, Supplementary Fig. 1**). Both manipulations induced a range of statistically significant transcriptional changes within OPCs compared to their respective controls (**Fig. 3b)**. Overall, we found that gene expression changes following demyelination were more pronounced than after 4AP treatment, with approximately twice as many significantly changed genes in demyelination than after activity stimulation (**Fig. 3b, c, Supplementary Table 1**). Interestingly, comparison of the identity of genes changed revealed that there was only minimal overlap with no more than 5% of differentially regulated genes being shared between conditions (**Fig. 3c**). Furthermore, the analysis of gene ontology (GO) terms to classify the differentially regulated genes into functional categories of biological processes revealed the activation of divergent transcriptional programmes between conditions. Major pathways altered in 4AP-mediated activity stimulation related to downstream ’signalling’, ’protein catabolism’, and ’regulation of neurite growth and motility’, accounting for 53 out of 218 (24%) differentially regulated genes (**Fig. 3d, Supplementary Table 1**). In contrast, major pathways activated in OPCs after demyelination related to cell ’metabolism’, ’extracellular matrix’ and ’differentiation’, accounting for 61 out of 414 (15%) differentially regulated genes (**Fig. 3e, Supplementary Table 1**). Closer analysis revealed that only in response to demyelination did we find the direct upregulation of genes associated with oligodendrocyte differentiation. In fact, the most significant of all 404 differentially regulated genes was *flj13639* (**Fig. 3b**). This gene encodes for a chain dehydrogenase called 36k which has important functions in regulating oligodendrocyte differentiation and is an abundant component of zebrafish myelin ^28,29^. Another significantly upregulated gene with direct links to oligodendrocyte biology was *gpr17* (**Fig. 3b**). Gpr17 is a G-protein coupled receptor that is specific to the oligodendrocyte lineage in the CNS. Single cell transcriptome analyses from human, mouse and zebrafish localise *gpr17* to clusters related to committed OPCs and pre-myelinating oligodendrocytes ^14,30,31^, where signalling via this receptor is believed to act as a break on differentiation in mammalian and zebrafish models ^32–35^. However, functional experiments using *in vivo* fate tracking provide a less clear picture on the identity and roles of Gpr17-expressing cells, depending on experimental context. For example, it was shown that the Gpr17-positive OPCs do not significantly contribute to the generation of myelinating oligodendrocytes in the healthy adult CNS, but that they can do so in response to CNS injury ^15,21,36^. Hence, we reasoned that studying differential upregulation of Gpr17 in response to enhanced motor activity and demyelination observed in our study might reveal insights into context-dependent mechanisms of oligodendrocyte recruitment.

**Figure 3:**
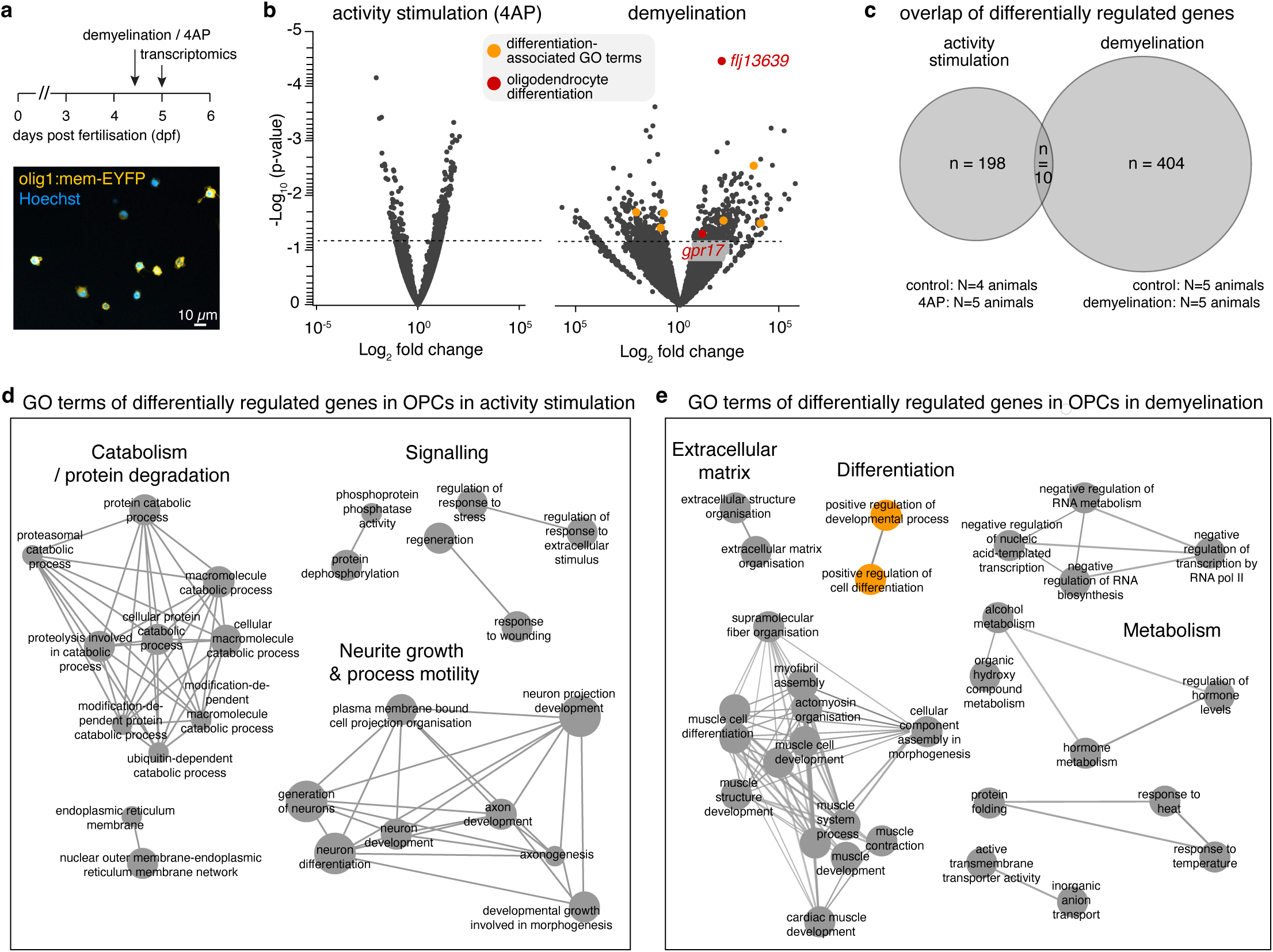
Divergent transcriptional programmes in OPCs after demyelination and activity stimulation. **a)** Schematic showing treatment paradigm used prior to fluorescence activated cell sorting of OPCs from Tg(olig1:mem-EYFP). Image shows example of sorted cells used for RNA extraction. Scale bar: 10µm. **b)** Volcano plots of gene expression changes in demyelination vs control or 4AP vs control. Data are plotted with log2 fold change on the x-axis, and -log10 p-value on the y-axis. Dotted line indicates significance threshold of p=0.05. Red and orange dots depict genes linked to differentiation. In the demyelination experiments, n=5 control and n=5 demyelination RNA samples were analysed. In the 4AP-treatment experiments, n=4 control and n=5 4AP RNA samples were analysed. Each sample was generated in a different experiment. **c)** Venn diagram showing the number of significantly changed genes that are shared between the demyelination and activity stimulation (4AP) groups. **d)** GO term analysis of the significantly differentially regulated genes in OPCs after activity stimulation. **e)** GO term analysis of the significantly differentially regulated genes in OPCs after demyelination (dfferentiation-linked terms highlighted in yellow).

### Live cell imaging of fluorescent knock-in reporter reveals properties, origins and fates of Gpr17-expressing oligodendrocyte lineage cells

To study the dynamics of Gpr17-expressing OPCs by live cell imaging we generated a novel fluorescent knock-in (KI) reporter line. In this line, EYFP followed by a P2A self-cleavage peptide was inserted directly after the start codon of the *gpr17* locus (**Fig. 4a, Supplementary Fig. 2**). This design allows for co-expression of the fluorescent reporter alongside the endogenous gene as separate proteins from a single mRNA as shown in previous studies ^27,37^. Crossing the EYFP-2A-Gpr17 KI reporter in a transgenic background that labels all OPCs (Tg(olig1:nls-Cerulean)) and myelinating oligodendrocytes (Tg(mbp:KR)) revealed that Gpr17-expressing cells were localised across the spinal cord where they could co-express either of the two other markers (**Fig. 4b**).

**Figure 4:**
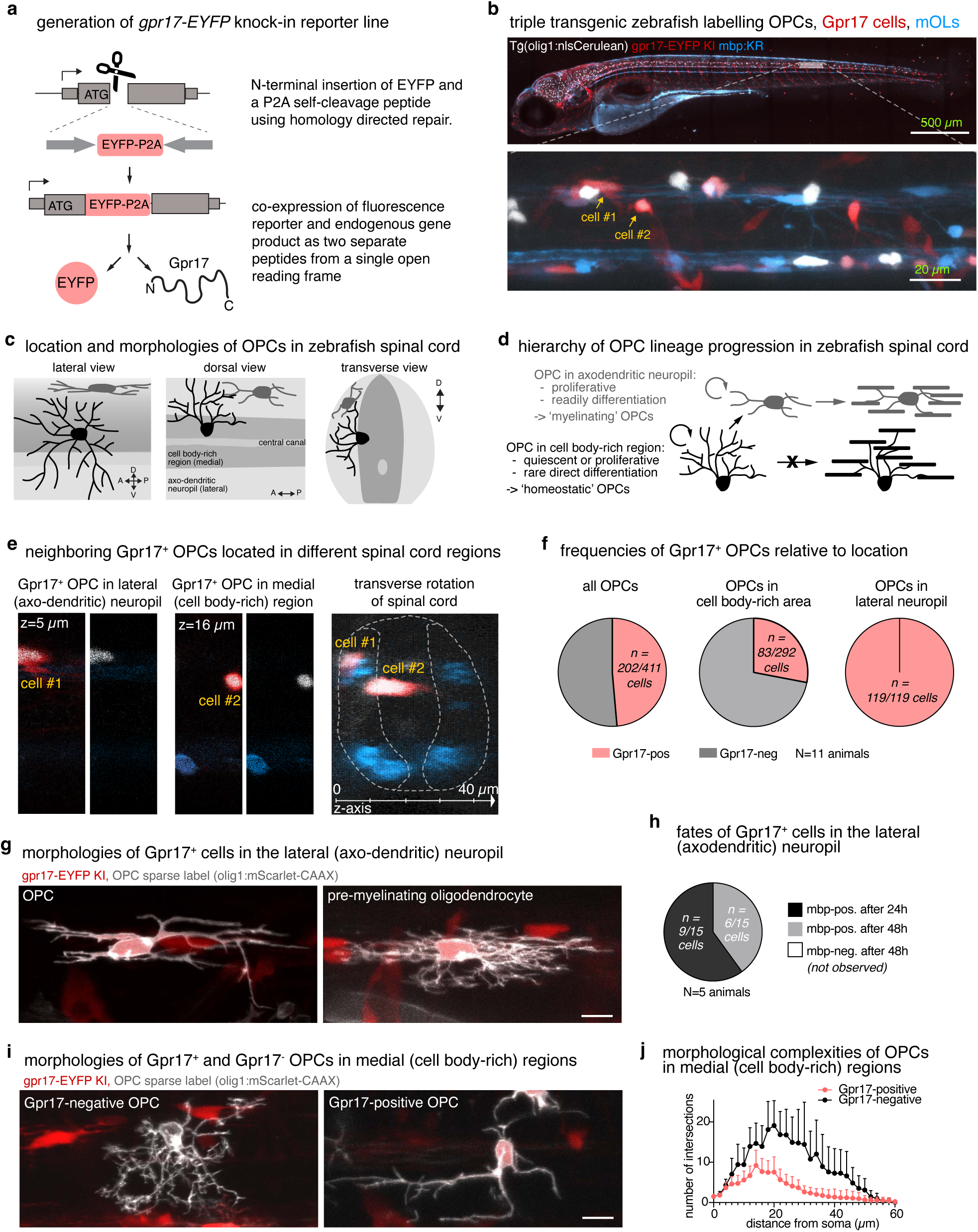
A subpopulation of OPCs identified by Gpr17 expression. **a)** Schematic depicting the strategy to generate the EYFP-P2A-gpr17 knock-in reporter line. **b)** Top: Overview image of a tripe transgenic zebrafish expressing Tg(olig1:nlsCerulean), EYFP-P2A-gpr17, and Tg(mbp:KillerRed). Scale bar: 500µm. Bottom: Zoom-in from the boxed area above showing a higher magnification view in a region of the spinal cord. Arrows pointing at ‘cell #1’ and ‘cell#2’ depict two closely adjacent gpr17-positive OPCs analysed in panel e. Scale bar, 20µm. **c)** Schematic summarising the location and morphologies of OPC in the zebrafish spinal cord. In brief, all OPCs extend their processes into the axo-dendritic neuropil of the lateral spinal cord. However, OPC somata can reside in the cell body-rich region of the medial spinal cord, or within the axo-dendritic neuropil of the lateral spinal cord. See ref. ^14^ for details. **d)** Schematic summarising the hierarchy of OPC lineage progression in the zebrafish spinal cord. In brief, ‘homeostatic’ OPCs in the cell-body rich region rarely directly differentiate, but they can divide to give rise to ‘myelinating’ OPCs which may migrate to the axo-dendritic neuropil, where they continue to proliferation, and ultimately differentiate. See ref. ^14^ for details. **e)** Example images of two neighbouring Gpr17-positive OPCs residing with their soma in the cell body rich and axo-dendritic area of the lateral spinal cord, respectively. **f)** Quantification of the frequency of Gpr17-positive OPCs in different areas of the spinal cord at 5 d.p.f.. N=411 cells from n=11 animals. **g)** Sparse labelling using olig1:mScarlet-CAAX to showing example morphologies of Gpr17-positive cells in the lateral axo-dendritic neuropil at 5 d.p.f. Scale bar, 10µm. **h)** Quantification of differentiation fates acquired by individual Gpr17-positive/mbp-negative in the axo-dendritic neuropil over time between 5-7 d.p.f.. N=15 cells from 5 animals. **i)** Sparse labelling using olig1:mScarlet-CAAX to showing example morphologies of Gpr17-negative and Gpr17-positive OPCs cell body-rich area of the medial spinal cord, respectively. Scale bar, 10µm. **j)** Sholl analysis to quantify ranching complexity of individual Gpr17-negative -positive OPCs in the cell body-rich area. Data shown as mean + 95% C.I.. N=15 Gpr17+ OPCs/n=12 Gpr17- OPCs.

Focussing on OPCs (i.e. olig1:nls-Cerulean-positive/mbp:KR-negative), we have previously carried out an in-depth characterisation of their properties and functions in the zebrafish spinal cord, where we established a lineage hierarchy within the OPC population (**Fig. 4c, d**) ^14^. In brief, the zebrafish spinal cord can be divided into two core regions: a cell-body rich area in the medial spinal cord and an axo-dendritic neuropil in the lateral spinal cord. OPCs could reside with their cell bodies in either of these regions while the processes of all OPCs always extend in the axo-dendritic neuropil (**Fig. 4c**). Clonal fate tracking revealed that OPCs could proliferate in either location. However, only the OPCs located in the axo-dendritic neuropil and not the ones in the cell body-rich area differentiated readily. Furthermore, this pool of OPCs was replenished by proliferation of OPCs in the medial, cell body-rich regions, followed by their subsequent migration to the axo-dendritic neuropil (**Fig. 4d**) ^14^. Hence, we termed the OPCs in the cell-body rich area ’homeostatic OPCs’ and the ones in the axo-dendritic neuropil ’myelinating OPCs’. Here, using our new reporters we saw that both subpopulations could express Gpr17 (**Fig. 4e**). Quantification revealed that approximately half of the entire OPC population expressed Gpr17, with 100% of the ’myelinating’ OPCs in the axo-dendritic neuropil and about one quarter of the ’homeostatic’ OPCs in the cell body-rich regions being Gpr17-positive (**Fig. 4f**). Sparse labelling of individual OPCs using a membrane-tethered fluorescent reporter (olig1:mScarlet-CAAX) showed that the Gpr17-labelled cells in the axo-dendritic neuropil had morphologies that were either reminiscent of OPCs (bearing exploratory processes only), or of pre-myelinating oligodendrocytes (being highly branched and likely forming nascent axonal contacts) (**Fig. 4g**). Fate tracking revealed that all Gpr17-expressing cells in the axo-dendritic neuropil upregulated myelin basic protein (mbp) reporters within 48 hours of analysis (**Fig. 4h**), confirming our previous observations on the fates of this OPC subpopulation in the zebrafish spinal cord, and confirming published literature on the identity of Gpr17-postive cells as committed precursors and pre-myelinating oligodendrocytes. However, we were surprised to see that 25% of the OPCs in the cell-body rich regions also expressed Gpr17, as we have previously shown that these cells rarely directly differentiate into myelinating oligodendrocytes ^14^ . Furthermore, analysis of cell morphology revealed that the Gpr17-positive OPCs in these regions exhibit much simpler process branching than their Gpr17-negative siblings (**Fig. 4i, j**). In fact, the morphology of the Gpr17-positive cells in the cell-body rich area was much more reminiscent of very immature and migratory OPCs, rather than OPCs that are primed to differentiate (as Gpr17 expression would suggest).

To gain further insight on the origins and fates of these immaturely appearing Gpr17-positive OPCs in the cell-body rich area, we carried out high-resolution timelapse imaging of our triple transgenic animals, sampling OPCs every 20 minutes for up to 72 hours starting at 3 days post fertilisation (**Fig. 5, Supplementary Movie 3**). Across all timelapses we observed the appearance of 40 newly formed Gpr17-expressing OPCs in the cell body-rich area of the medial spinal cord. All these cells (n=40/40) were associated with a cell division (**Fig. 5c chart #1**). Of these proliferation-associated events, 40% (n=16/40) emerged from a Gpr17-negative parent cell and the other 60% (n=24/40) came from a division of a Gpr17-positive parent cell in the cell body-rich area (**Fig. 5c chart #2**). Not all divisions of Gpr17-negative OPCs gave rise to Gpr17-positive daughter cells (n=18/28 divisions observed, **Fig. 5c chart #3**). Interestingly, however, fates of divisions were always symmetric in that either both (n=18) or none (n=10) of the daughter cells expressed Gpr17 (**Fig. 5c chart #4**). We did not observe a Gpr17-positive OPC giving rise to Gpr17-negative cells. Once formed, about three quarters of newly formed Gpr17-positive OPCs in the cell body-rich areas migrated to the axo-dendritic neuropil over the course of our analysis (n=32/41, **Fig. 5d**), where we continued to observe cell divisions in 7/32 occasions (**Fig. 5e**). These observations are consistent with the simple, migratory morphology of these Gpr17-positive OPCs (**Fig. 4i, j**). Quantification of the timings of Gpr17 upregulation, migration to lateral spinal cord, and subsequent differentiation showed that these events followed a stereotyped temporal pattern for all cells studied (**Fig. 5f**). Therefore, we conclude that Gpr17-positive OPCs emerge from symmetrically-fated proliferation events of Gpr17-negative OPCs in the cell body-rich area, and by proliferation-mediated expansion of Gpr17-positive cells (**Fig. 5g)**.

**Figure 5:**
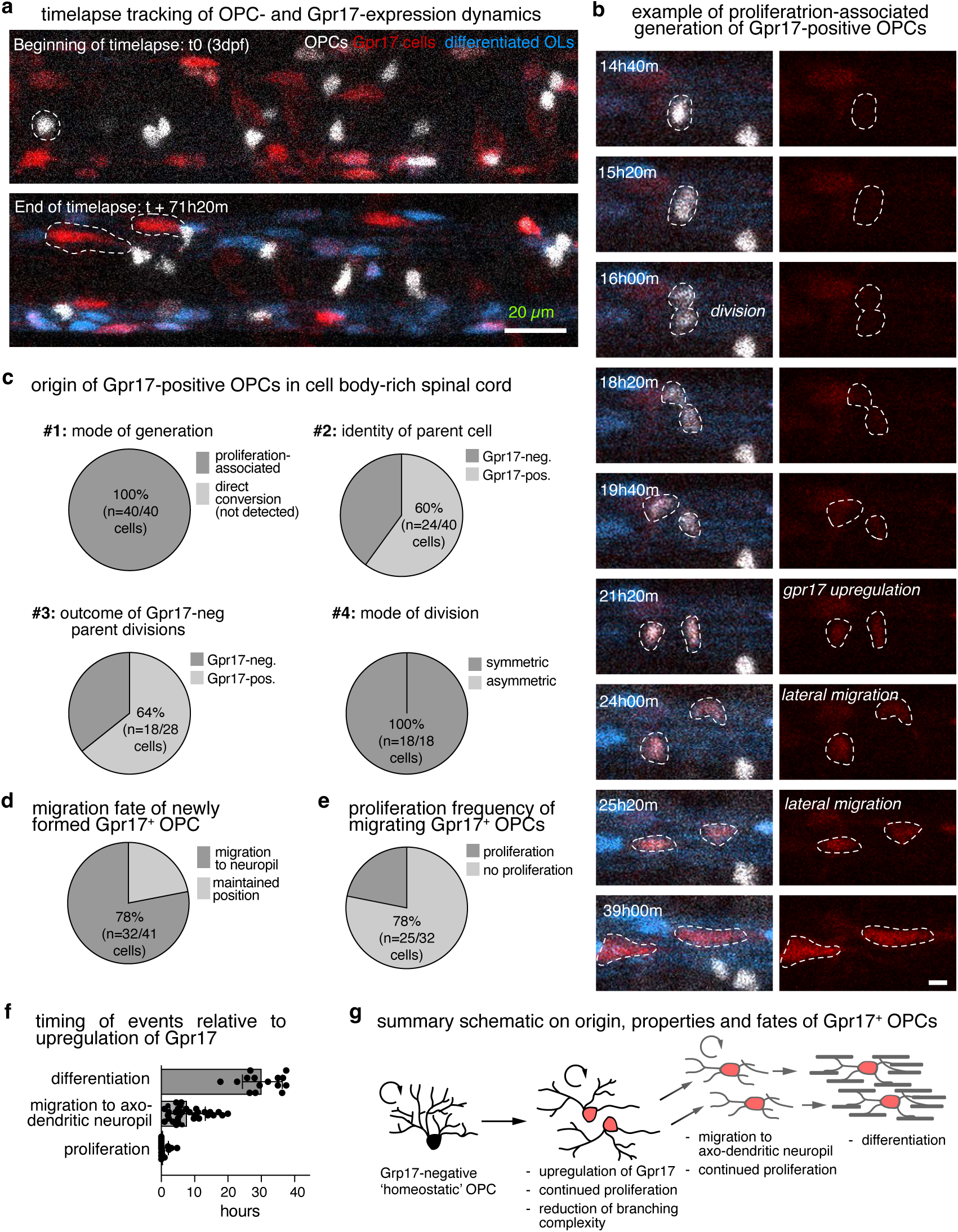
New Gpr17-expressing OPCs emerge from symmetric divisions. **a)** Example images of the first (top) and last (bottom) frame of a three day time lapse in the spinal cord of a Tg(olig1:nlsCerulean), Tg(mbp:KillerRed), Gpr17-EYFP animal between 3-6 d.p.f. Arrows depict example cells shown in panel b. Scale bar, 20µm. **b)** Example of individual Gpr17-negative OPC followed over time as it produces Gpr17-positive daughter cells. Scale bar: 5µm. **c)** Pie charts showing frequencies of observations on the origin of new Gpr17-positive OPCs in the medial spinal cord. Data were collected from three time lapse recordings of Tg(olig1:nlsCerulean), EYFP-P2A-Gpr17, Tg(mbp:KillerRed) animals starting at 3 d.p.f.. N=3 animals. **d)** Pie chart showing frequency of migration fates of newly formed Gpr17-positive OPCs in the medial spinal cord OPCs within 24h. N=3 animals. **e)** Pie chart showing proliferation frequency of Gpr17-posive OPCs after they migrated to the lateral spinal cord. N=3 animals. **f)** Quantification of the timing of proliferation, lateral migration to axo-dendritic area, and differentiation relative to Gpr17-upregulation (indicated by dotted line). N=18/32/15 cells from 3 animals. **g)** Summary schematic on the origin, properties and fates of Gpr17+ OPCs. In brief, most new Gpr17-positive OPCs arise from symmetric divisions of Gpr17- parent cells (homeostatic OPCs). Newly formed Gpr17-positive OPCs have a much simpler morphology than their parent cells and frequently migrate to the axo-dendritic neuropil where they can continue to proliferate and eventually differentiate.

### Demyelination unlocks a differential mode of Gpr17-expressing OPC production

We have previously shown that activity stimulation with 4AP initially enhances OPC proliferation, with a subsequent increase in differentiating oligodendrocyte numbers ^14^. Confirming these findings, we also saw increased OPC numbers in the axodendritic neuropil with 4AP treatment in the present study (**Supplementary Fig. 3**). However, this was not the case after demyelination **Supplementary Fig. 3a**), indicating the presence of differential routes of oligodendrocyte recruitment.

Having shown in our triple marker analyses that all newly generated myelinating oligodendrocytes emerge from a Gpr17-expressing state, and that new Gpr17-positive OPCs exclusively associated to cell divisions, we wanted to test if demyelination and activity-stimulation give rise to Gpr17-positive OPCs in a differential manner. To do so, we focussed on the presence of Gpr17-positive OPCs located in the cell body-rich area of the medial spinal cord as a simple readout, because this is where the new OPCs fated for myelination emerge first (**Figs. 4, 5**). In these regions, we observed a ∼50% increase in their numbers at 1 day post demyelination (**Fig. 6a**). By contrast, neither of our two independent assays of enhanced motor activity showed this change when compared to their respective controls (**Fig. 6b, c**). This observation suggests that specifically in response to demyelination, a higher proportion of Grp17-positive OPCs are generated in this region, parallelling our RNA sequencing results. To test if these are generated via the same default mode where their production is directly linked to cell divisions, we clonally fate-tracked individual Gpr17-negative OPCs in the cell body-rich area through demyelination. To our surprise, following demyelination we frequently observed the direct upregulation of Gpr17 that was not linked to a division event (**Fig. 6d**). We also noted that the morphology of pre-existing individual OPCs transformed from highly complex prior to demyelination (characteristic of homeostatic OPCs), to a much simpler morphology with fewer and longer processes as the cell upregulated the Gpr17 reporter (characteristic of migratory OPCs, **Fig. 6d**). This further supports our previous reports on the morphology of homeostatic OPCs typically seen in cell body rich areas ^14^, and the divergent, migratory morphology of the Gpr17-positive OPCs in the same regions that we report here (**Fig. 4h, i**). When we quantified the frequency by which this newly observed mode of Gpr17-positive OPC generation occurred in demyelination, our clonal analyses revealed that new Gpr17-positive OPCs emerged from a negative parent cell through direct conversion in about 1/3 of all cases, with the remainder being proliferation-associated events (**Fig. 6e**). By contrast, in controls all new Gpr17-positive OPCs derived from proliferation-associated events, as established earlier (**Fig. 6e**, **Fig 4**.). Hence, we conclude that demyelination but not activity stimulation unlocks an additional mode of oligodendrocyte recruitment from the pool of OPCs.

**Figure 6:**
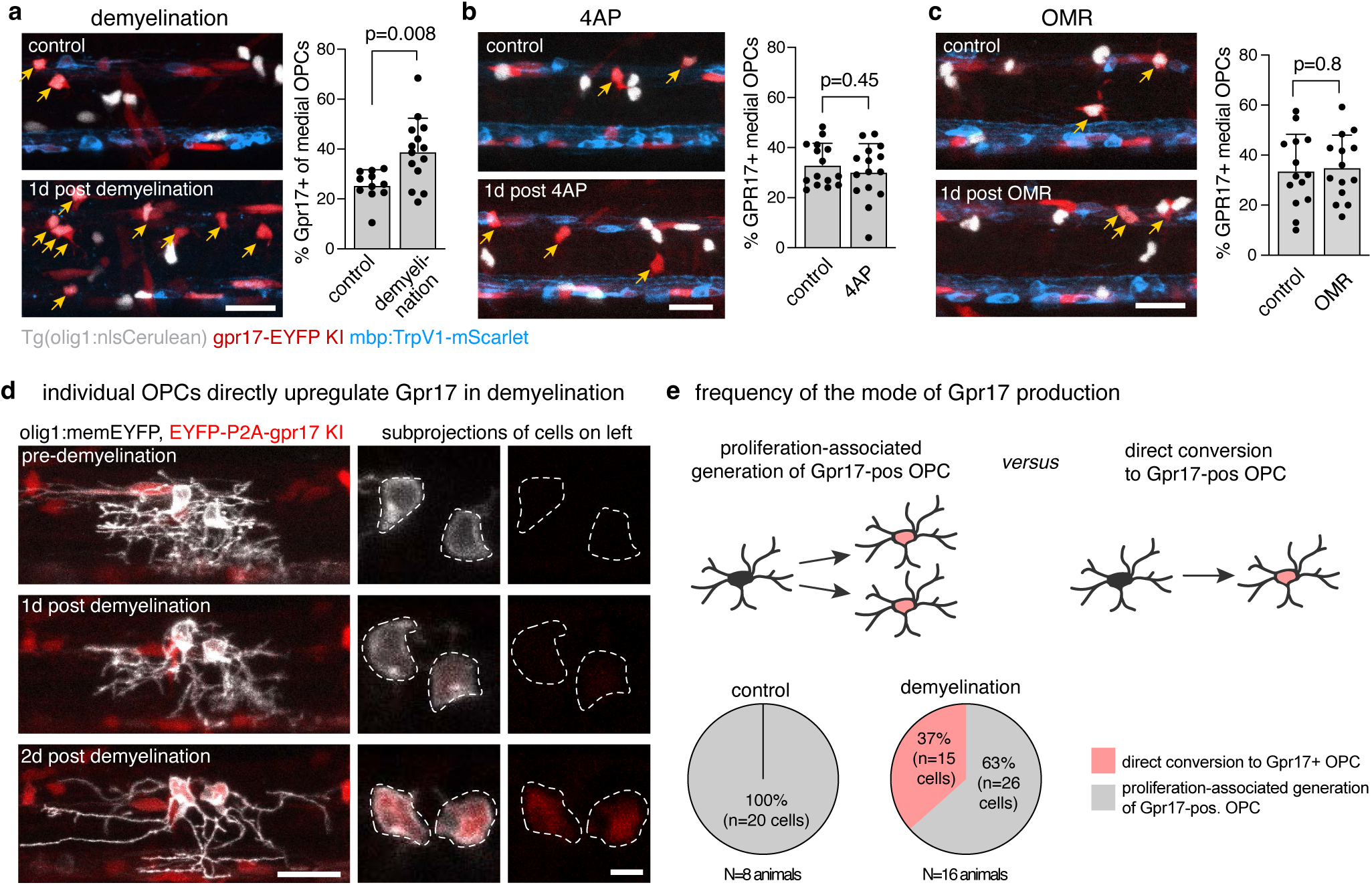
Proliferation independent, direct conversion to Gpr17-positive OPCs occurs selectively in demyelination. **a)** Example images of triple transgenic animal 1d post demyelination at 5 d.p.f.. Yellow arrows indicate presence of Gpr17-positive OPCs in the cell body rich areas. Scale bar, 20µm. Data in the graph shows the respective quantification, and is shown as mean ± S.D % medial Gpr17-positive OPCs in control vs demyelination (25.6±5.8 *vs* 39.0±13.9; p=0.008, Mann Whitney test, *U* = 29.5. N=11/14 animals in control/demyelination. **b)** Example images as in panel a, here comparing the effects of 4AP on Gpr17-positive OPCs in the cell body rich areas (yellow arrows). Scale bar, 20µm. Data in the graph shows the respective quantification, and is shown as mean ± S.D % medial Gpr17-positive OPCs in control vs 4AP (33.1±8.4 *vs* 30.3±11.2. p=0.45, unpaired two-tailed t test, t=0.775, df=28). N=15/15 animals in control/4AP. **c)** Example images as in panel a, here comparing the effects of OMR on Gpr17-positive OPCs in the cell body rich areas (yellow arrows). Scale bar, 20µm. Data in the graph shows the respective quantification, and is shown as mean ± S.D. % medial Gpr17-positive OPCs in control vs OMR (33.8±14.9 vs 35.1±12.6, p=0.8, unpaired two-tailed t test, t=0.26, df=26). N=14/14animals in control/OMR. **d)** Example images of 2 individual olig1:memEYFP labelled OPCs in Gpr17-EYFP KI zebrafish, before and after demyelination (4 d.p.f. – 6 d.p.f.). Note the reduction in process complexity as the cell begins to express Gpr17. Scale bars: 20µm (overview), 5µm (zoom-in). **e)** Top: Schematic of the potential modes by which new Gpr17+ OPCs can be generated from a Gpr17 negative OPC. Bottom: pie charts showing the frequency of the mode of new Gpr17+ OPC production in control and after demyelination. N=20/41 cells in 8/16 animals in control/demyelination.

## Discussion

In this study we have established independent assays to increase oligodendrocyte generation in response to enhanced motor activity and demyelination in zebrafish, respectively. Our rationale for these experiments was that it allows studying these processes using high-resolution *in vivo* fate tracking in a relatively simple model system, where most experimental parameters remain the same apart from the upstream stimulus to trigger enhanced oligodendrogenesis. We reasoned that this would allow us to directly test for context-dependent differences in the mechanisms of oligodendrocyte recruitment from the OPC pool. Our experiments provide novel insights in the gene expression changes of OPCs in response to enhanced activity and demyelination, the identity and properties of the OPC subpopulation labelled by Gpr17, and on the mechanisms of oligodendrocyte recruitment in plasticity and repair, respectively.

Even though activity and demyelination both enhanced the rate of oligodendrocyte formation, our transcriptomic analyses of isolated OPCs showed that they undergo highly divergent transcriptional changes in response to our manipulations. On the one hand, this is not unexpected because the upstream stimuli are quite different. For example, a hallmark of demyelination is the activation and influx of phagocytes which are known to release a host of molecules that alter inflammatory and regenerative responses. In contrast, a key feature of the 4AP-mediated activity stimulation is enhanced calcium signalling activity in neurons and OPCs without inflammation. However, it was a novel insight that only demyelination resulted in the direct upregulation of differentiation-associated genes, even though we and others have previously shown that manipulation of activity in general, and of calcium signalling in particular, regulates oligodendrogenesis ^38,39^ . We can only speculate which of these transcriptional changes lead to increased myelinating oligodendrocyte numbers when there is no direct increase in differentiation genes. It is possible that activity manipulations primarily act on other aspects of the oligodendrocyte lineage, where increased myelination is a secondary outcome of multiple changes. For example, we have previously shown that the most immediate effect of 4AP is on OPC proliferation ^14^. Furthermore, a very recent study showed that enhanced neuronal activity increases the successful integration of differentiating OPCs/pre-myelinating oligodendrocytes ^40^. Therefore, it is possible that enhanced myelination can occur without directly increasing OPC differentiation, but rather by *i)* increasing numbers of differentiation-primed OPCs, and *ii)* by better maintaining those when they go on differentiate. Such a model would be consistent with the data presented in our study, which suggest that demyelination but not enhanced activity lead to a direct increase in OPCs entering a differentiation programme.

We were particularly intrigued to see that the marker Gpr17 was selectively upregulated in our transcriptomic data following demyelination but not enhanced motor activity. On the one hand Gpr17 is tightly linked to oligodendrocyte differentiation in single cell transcriptomics where it marks clusters of pre-myelinating oligodendrocytes and OPCs committed/primed to differentiate ^32–35^. On the other hand, fate tracking and genetic manipulation studies indicate that pathways of oligodendrocyte differentiation exist that do not involve Gpr17, and that these may differ under physiological and pathological conditions ^15,21,36^. Hence, we generated a fluorescent Gpr17 KI reporter to elucidate whether OPC fate tracking alongside that of Gpr17 expression may reveal different modes of oligodendrocyte generation. We see that in our model all new myelinating oligodendrocytes come from Gpr17-positive cell states, supporting the single cell transcriptomic data. However, our reports do not exclude the possibility that alternative oligodendrocyte differentiation may exist in zebrafish as reported in mice, for example at later developmental stages / adulthood. Our fate tracking further revealed that in healthy conditions, all new Gpr17-expressing OPCs emerged from proliferation-associated events and that these divisions were always symmetrically fated, *i.e.* either both or none of the daughter cells upregulated Gpr17. Although we have no explanation for the underlying reasons, our observations suggest that the decision whether to upregulate Gpr17 expression (*i.e.* priming them towards differentiation) is likely determined before the division occurs. Consequently, OPCs may exhibit two different types of divisions: one that is for maintenance of the OPC pool, and one that is to produce OPCs that will go on to differentiation.

While the link between recent OPC division and subsequent differentiation is strong during earlier phases of development in zebrafish and mice ^14,16^, it is less clear in adulthood where direct differentiation of OPCs that cannot be linked to a recent division has been observed ^18,20^. Future studies will be required to directly address if the mode of oligodendrocyte generation changes from development to adulthood. Alternatively, different and/or additional OPCs with different properties may exist in the adult but not in the young animal. Indeed, electrophysiological and transcriptomic studies have shown that OPCs increasingly diversify with age ^13^. However, markers to differentiate between OPCs with different properties are notoriously lacking; Gpr17 being one of very few that has been assigned to label a subpopulation of OPCs. Interestingly, in our work we observed that even though Gpr17 expression is highly predictive of future differentiation, the morphology of newly formed Gpr17-positive OPCs was much more reminiscent of a very immature OPC, even when compared to their Gpr17-negative parent cells. This has important implications as it means that cell morphology alone can be deceptive in drawing conclusions on future fates of an OPC.

A key finding from the work presented here is that the mechanisms of oligodendrocyte recruitment from the OPC pool are different in plasticity and repair. Importantly, only the tracking of individual OPCs whilst monitoring their Gpr17-expression state revealed this direct recruitment of Gpr17-positive OPCs from the pool of homeostatic OPCs that we only observed in demyelination.

Therefore, we conclude that regenerative myelination unlocks an additional mode of oligodendrocyte recruitment, whereas adaptive/ activity-induced myelination may rather enhance the gain of a default mode of oligodendrocyte recruitment. Our result provides an answer to a long-standing question in the field as to whether remyelination is simply a recapitulation of development^41^. However, several questions remain to be addressed in future studies. For example, how do the findings reported here compare to other age-related of physiological contexts where the pool of OPCs may be different? Another key remaining task will be to elucidate the signalling pathways that lead to the direct transition to a Gpr17-positive state, as the upregulation of this receptor reflects an outcome of (yet unknown) signalling that likely occurred at earlier stages in cells that we consider as homeostatic OPCs in our model. Identifying these triggers will be a step towards being able to steer oligodendrocyte lineage progression in contexts where these processes are inefficient.

## Acknowledgements

We thank Wenke Barkey for excellent technical support, Julia Meng for help with the optical treadmill code, David Lyons for supervision of some of the optical treadmill assays, and all members of the Czopka group for feedback on the manuscript. This work was funded by grants from the European Research Council (Horizon 2020 ERC StG grant MecMy, 714440) and the BBSRC (BB/V017012/1 and UKRI1946) to TC. KB was funded by the MRC doctoral training programme ’Precision Medicine’.

## Author contributions

Conceptualization: TC; Methodology: LJH, PB, EC, FA; Investigation: LJH, PB, EC, KB; Formal analysis: LJH, PB, EC; Supervision: TC; Visualisation: LJH, TC; Writing: TC; Funding acquisition: TC.

---

**Supplementary Figure 1:**
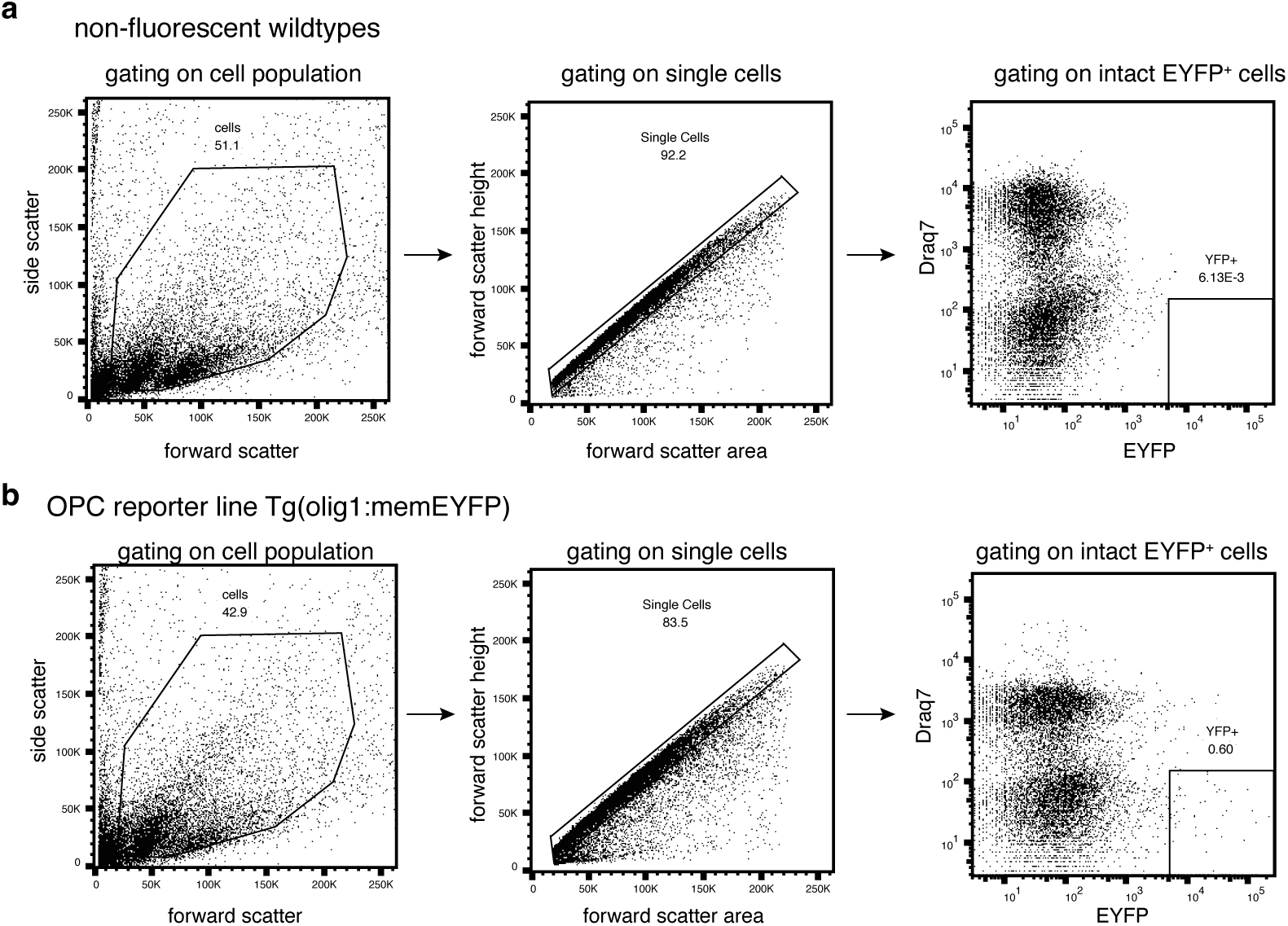
Fluorescent-activated cell sorting (FACS) of OPCs. Example plots showing the gating strategy to isolate intact single EYFP-positive cells from non-fluorescent wildtype controls (panel **a**) and from Tg(olig1:memEYFP) (panel **b**).

**Supplementary Figure 2:**
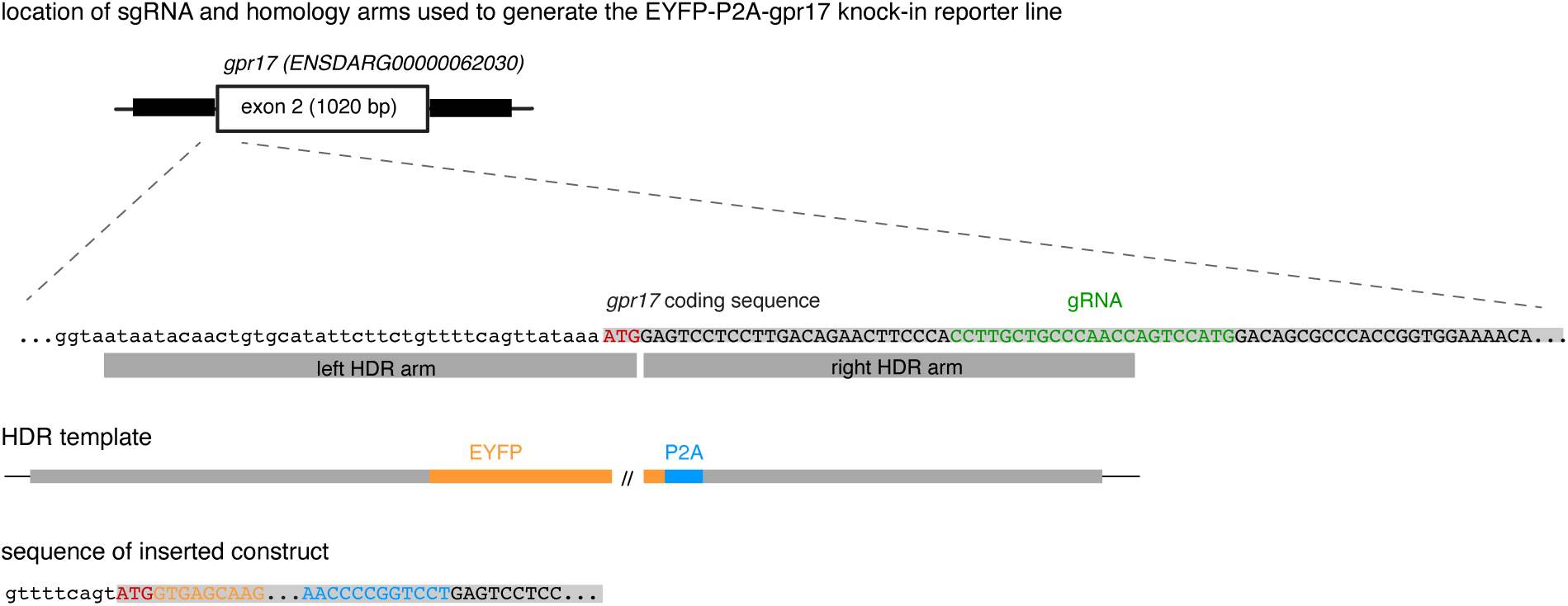
Generation of EYFP-P2A-gpr17 knock-in reporter line. Schematic cartoon summarising the location and sequences of key elements used for the generation of the EYFP-P2A-gpr17 knock-in reporter line. See also Supplementary Table 2 for details.

**Supplementary Figure 3:**
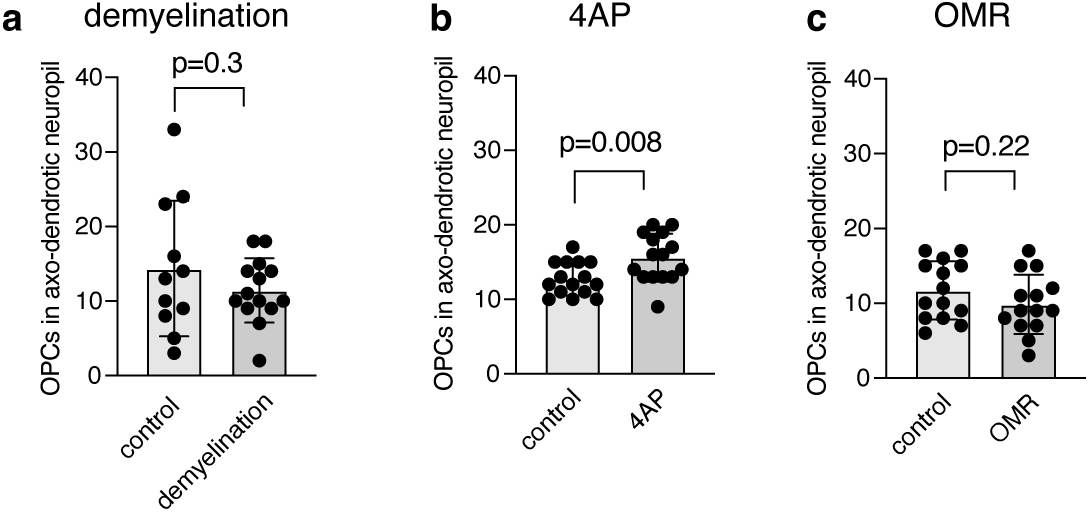
OPC numbers in axodendritic neuropil after demyelination and activity manipulations. **a)** Quantification of OPC numbers in axodendritic neuropil in control and 1 day post demyelination. Data are shown as mean ± S.D % lateral Gpr17-positive OPCs in control vs demyelination (14.4 ± 9.0 *vs* 11.4 ± 4.2; p=0.3, unpaired two-tailed, t=1.07, df=23). N=11/14 animals in control/demyelination. **b)** Quantification of OPC numbers in axodendritic neuropil in control and 1 day post 4AP treatment. Data are shown as mean ± S.D % lateral Gpr17-positive OPCs in control vs 4AP (12.7 ± 2.2 *vs* 15.6 ± 3.2. p=0.008, unpaired two-tailed t test, t=2.84, df=28). N=15/15 animals in control/4AP. **c)** Quantification of OPC numbers in axodendritic neuropil in control and 1 day post 4AP treatment. Data are shown as mean ± S.D % lateral Gpr17-positive OPCs in control vs OMR (11.7 ± 2.9 vs 9.9 ± 3.8, p=0.22, unpaired two-tailed t test, t=1.25, df=26). N=14/14animals in control/OMR.

**Supplementary Table 1: List of differentially regulated genes in OPCs after 4AP treatment and demyelination, respectively.**

**Supplementary Table 2: List of primers used**

**Supplementary Movie 1: Zebrafish motor behaviour elicited by optic treadmill**

**Supplementary Movie 2: Zebrafish motor behaviour elicited by 4AP treatment**

**Supplementary Movie 3: Timelapse analysis OPC dynamics, Gpr17-expresion, and differentiation fate in triple transgenic reporter line**

## Methods Details

### Zebrafish lines and husbandry

We used the following existing zebrafish lines and strains: Tg(mbp:nls-EGFP)^zf3078Tg^ ^42^ , Tg(mbp:KillerRed)^tum103Tg^ ^14^ , Tg(mfap4:memCerulean)^tum104tg^ ^14^ Tg(mbp:EGFP-CAAX)^ue2Tg^ , ^14^Tg(olig1:memEYFP)^tum107Tg^ ^14^, Tg(olig1:nls-Cerulean)^tum108Tg^ ^14^, and corresponding wildtypes in an AB or nacre/TL background. The following lines were newly generated for this study: Tg(mbp:TrpV1-mScarlet), Tg(mbp:mScarlet-CAAX), EYFP-P2A-gpr17 knock in. All animals were kept at 28.5 degrees with a 14/10 hour (h) light/dark cycle according to local animal welfare regulations. All experiments carried out with zebrafish at protected stages were approved by the governments of Upper Bavaria (animal protocols AZ55.2-1-54-2532-199-2015 and AZ55.2-1-54-2532-200-2015 to T.C.) and the UK Home Office (project licenses PP4944080 to T Czopka and PP3290955 to D. Lyons).

### Transgenesis constructs and generation of transgenic lines

We used the following existing expression constructs: pTol2_olig1(4.2):mScarlet-CAAX ^43^, pTol2_olig1(4.2):memEYFP ^44^. The following expression constructs were newly generated for this study: pTol2_mbp:TrpV1-mScarlet, pTol2_mbp:mScarlet-CAAX. To obtain these constructs, multi-site LR recombination reactions were performed using the following clones p5E_mbp ^45^ , pME_mScarlet-CAAX ^43^ , pME_TrpV1_nostop (newly generated for this study), p3E_mScarlet_pA ^46^ , and components of the Tol2kit ^47^ . pME_TrpV1_nostop was generated by PCR amplifying the coding sequence from a template plasmid using the primers attb1-TrpV1 and attB2R-TrpV1 (**Supplementary Table 2**) and subsequent BP recombination reaction with a pDONR221 vector. The plasmid pCS2+_EYFP-P2A was generated to serve as a PCR template for generation of homology-directed repair templates for making the EYFP-P2A Gpr17 knock-in line. Fertilised zebrafish eggs were microinjected with 1nl of an injection solution containing 5-25ng/µl DNA, 25-50ng/µl Tol2 transposase mRNA and 10% phenol red. Injected F0 animals were either used for single cell analysis, or raised to adulthood to generate full transgenic lines. For this, adult F0 animals were outcrossed with wildtype zebrafish and F1 offspring was screened for germline transmission of the fluorescent transgene.

### CRISPR/Cas9-mediated generation of Gpr17 knock-in reporter line

A knock-in line where EYFP-P2A is knocked-in at the start codon of the *gpr17* gene was established using CRISPR/Cas9-mediated insertion of a PCR-generated homology-directed repair (HDR) template using a previously described method ^48^ . Sequences for all primers used are listed in **Supplementary Table 2**. Primers were used to generate a gRNA targeting Gpr17 exon 1 using the GeneArt Precision gRNA synthesis kit (ThermoFisher). An HDR template, consisting of EYFP-P2A flanked by 40bp long homology arms targetting the desired integration side was generated by PCR using Q5 polymerase (New England Biolabs), and gel-purified using the Monarch PCR and DNA cleanup kit (New England Biolabs). Injection solutions were prepared in nuclease-free water to the final concentrations as follows: gRNA (12.5ng/µl), HDR template (100ng/µl), HDR modulator NU7441 (Stratech, 73μM), 10% phenol red and Cas9 enzyme (500ng/µl). F0 animals showing widespread fluorescence in oligodendrocyte-lineage cells were raised and outcrossed to wild types to establish stable F1 knock-in animals. The exact knock-in was confirmed by DNA sequencing and is as shown in **Supplementary Fig. 2**.

### Experimental manipulations to enhance oligodendrogenesis

Demyelination assays were performed by incubating Tg(mbp:TRPV1-mScarlet) larvae overnight in 20μM capsaicin (Sigma-Aldrich) dissolved in Danieau’s buffer. Larvae were kept in the dark during the incubation. The larvae were then washed twice in Danieau’s buffer before imaging was performed.

4AP incubation assays to study effects on oligodendrogenesis were performed by incubating larvae for 2 days in 0.1mM 4AP (Sigma Aldrich), dissolved in Danieau’s buffer. After the first day, the larvae were incubated in Danieau’s buffer for a 2h break for feeding, and then fresh 4AP was added.

For visually enhanced motor activity, *a* browser-based program called “Optical Treadmill” (https://github.com/klathem/Optical-Treadmill) was ran on an iPad to project moving gratings to 15 –50 zebrafish placed in a 100mm diameter Petri dish placed on the iPad screen inside an incubator heated to 28.5°C. The moving gratings consisted of alternating black and white rectangles with a 20-pixel width at a speed of 180 pixels/second. Chronic stimulation protocol was applied for 30 hours, and consisted of cycling a 30-minute loop of alternating 30-second right and left moving gratings followed by a 45-minute break where a black screen was shown. To enhance oligodendrogenesis, this assay was performed from 4-5 d.p.f. The control group was shown static gratings of the same size during the 30-minute stimulation blocks.

### Assessment of zebrafish swimming behaviour

Experiments were conducted on a custom-built, temperature-controlled behavioural rig set to 28.5C following build plans from the Stytra package ^49^. Briefly, behaviour was recorded using a high-speed camera (MQ013MG-ON, Ximea xiQ) fitted with a fixed focal length lens (#59-871, f = 25 mm, Edmund Optics) and an infrared long-pass filter (#66-106, 830 nm, Edmund Optics) to block visible light. The camera was positioned above the arena, which was illuminated from below using a 100 x 100 mm LED-based diffusive backlight (#66-839, 880 nm, Advanced Illumination). Visual stimuli were projected onto a diffusive screen placed 5 mm below the arena via a cold mirror (#62-633, Edmunds Optics) using a portable LED projector (P3E, Asus). Arenas were made by pouring 2% agarose (UltraPure Agarose #16500-500, Invitrogen by Life Technologies) in Danieau’s solution around a glass mold placed in a 90mm petri dish (Thermofisher). Each circular arena had a diameter of 50 mm with a graded slope towards a maximum depth of 5 mm. Fish position, heading direction and tail angles were tracked in real-time using the Stytra package in Python ^49^ . Fish with over 10% of missing tracking data were not considered for analysis. Distance swum was quantified using a custom-written Python script.

For assessment of 4AP treatment on motor activity, behaviour was recorded at 500 frames per second with a frame size of 640 x 512 pixels and an exposure time of 1.8 ms covering an area just larger than the arena, with a resolution of 0.05 mm per pixel. Individual larvae 5 dpf were recorded before and 25 minutes after being immersed in 0.1mM 4-AP dissolved in Danieau’s solution. Pre-and post-treatment, swimming behaviour was recorded for 3 minutes (1 minute habituation, 2 minutes exploration) while being presented with a blank, white screen.

For assessment of motor activity induced by the optical treadmill, behaviour was recorded at 200 frames per second with a frame size of 1280 x 1024 pixels and an exposure time of 1.8 ms covering an area just larger than the arena, with a resolution of 0.05 mm per pixel. Individual larvae 5-7dpf were placed in a circular arena filled with Danieau’s solution. Swimming behaviour was recorded for 35 minutes. First, a 5-minute habituation period during which a blank, white screen was presented to the animal. Then, each larva was presented with either a blank, white screen or moving gratings for 30 minutes. Gratings were 10mm in width and moving at a speed of 10 mm per second. Grating direction alternated between left and right every 30 seconds.

### Fluorescence activated cell sorting of single zebrafish OPCs and RNA isolation

Approximately 2000 Tg(olig1:memEYFP) at 5 dpf were euthanised and de-yolked by repetitive pipetting embryos in de-yolking buffer (55 mM NaCl, 1.2 mM KCl, 1.25 mM NaHCO_3_) with a P1000 pipette tip. Following two wash steps in Danieau’s buffer and centrifugation for 1 min at 300 g, tissues were digested for 30 minutes at 37°C in a shaking incubator using the Papain Dissociation Kit (Worthington Biochemical Corporations) according to manufacturer’s instructions. The obtained cell pellet was resuspended in FACSmax Cell Dissociation Buffer (Amsbio) with 5-10% FCS and filtered through a 30 µm Filcon syringe (BD Bioscience) before sorting. Approximately 100,000 EYFP^+^ / Draq7^-^ (Thermo Fisher Scientific) cells were sorted on an FACS Aria II cell sorter (Bectin Dickinson) to exclude dead cells and isolate a EYFP^+^ cell population. RNA was isolated from sorted EYFP+ cells using the RNeasy micro UCP kit (QIAGEN), as per kit instructions, and shipped on -80 to Qiagen for sequencing.

### RNA sequencing and analysis

Bulk RNA sequencing of isolated mRNA and bioinformatic analyses were carried out by QIAGEN as follows: "RNA integrity was assessed by Agilent TapeStation and only samples with RNA integrity values higher than 7 were used. The library preparation was done using the QIAseq UPX 3’™ Transcriptome Kit (QIAGEN). A total of 1ng or 4ng purified RNA was converted into cDNA NGS libraries. During reverse transcription, each cell is tagged with a unique ID and each RNA molecule is tagged with a unique molecular index (UMI). Then RNA was converted to cDNA. The cDNA was amplified, during the PCR indices were added and the libraries were purified. Library preparation was quality controlled using capillary electrophoresis (Agilent DNA 7500 Chip). Based on quality of the inserts and the concentration measurements the libraries were pooled in equimolar ratios. The library pool(s) were quantified using qPCR. The library pool(s) were then sequenced on a NextSeq (Illumina Inc.) sequencing instrument according to the manufacturer instructions with 100bp read length for read 1 and 27bp for read2. Raw data was de-multiplexed and FASTQ files for each sample were generated using the bcl2fastq2 software (Illumina inc.). The “Demultiplex QIAseq UPX 3’ reads” tool of the CLC Genomics Workbench 22.0.2 was used to demultiplex the raw sequencing reads according to the sample indices. The “Quantify QIAseq UPX 3’ workflow” was used to process the demultiplexed sequencing reads with default settings. In short, the reads are annotated with their UMI and are then trimmed for poly(A) and adapter sequences, minimum reads length (15 nucleotides), read quality, and ambiguous nucleotides (maximum of 2). They are then deduplicated using their UMI. Reads are grouped into UMI groups when they (1) start at the same position based on the end of the read to which the UMI is ligated (i.e., Read2 for paired data), (2) are from the same strand, and (3) have identical UMIs. Groups that contain only one read (singletons) are merged into non-singleton groups if the singleton’s UMI can be converted to a UMI of a non-singleton group by introducing an SNP (the biggest group is chosen). The reads were then mapped to the Zebrafish genome GRCz11 and annotated using the ENSEMBL GRCz11 v. 103 gene annotation.

The ‘Empirical analysis of DGE’ algorithm of the CLC Genomics Workbench 22.0.2 was used for differential expression analysis with default settings. It is an implementation of the ‘Exact Test’ for two-group comparisons developed by Robinson and Smyth, 2008 and incorporated in the EdgeR Bioconductor package Robinson et al., 2010.

For all unsupervised analysis, only genes were considered with at least 10 counts summed over all samples. A variance stabilizing transformation was performed on the raw count matrix using the function vst of the R package DESeq2 version 1.28.1. 500 genes with the highest variance were used for the principal component analysis. The variance was calculated agnostically to the pre-defined groups (blind=TRUE). 35 genes with the highest variance across samples were selected for hierarchical clustering."

### Gene Ontology term analysis

GO term analysis of up- and down- regulated genes was performed using the ClueGo plugin for Cytoscape ^50^. For the activity-stimulated group, we set the parameters as a miniumum of 2% of genes, and 6 genes minimum from the following databases: GO/Biological Processes (27.03.2019), GO/Molecular Function (27.03.2019) and Go/Cellular Component (27.03.2019). For the demyelination group, the parameters were set as a minimum of 3% of genes, and 5 genes minimum from the same databases. The kappa score was set to 0.4 for all analyses.

### Hybridisation Chain Reaction (HCR) for cfos detection

All reagents were ordered from Molecular Instruments and all HCR steps were performed in the dark. Briefly, whole larvae were sacrificed then fixed in 1mL of 4% paraformaldehyde overnight at 4°C, washed three times in phosphate-buffered saline (PBS) then dehydrated and rehydrated in successive 5 min MeOH/PBS-Tween (0.1%) washes (25%, 50%, 75%, 100%). The samples were incubated twice at 37°C with 100μL pre-warmed probe hybridization buffer for 15 min, then incubated overnight in 8nM of *cfos* probe diluted in probe hybridization buffer (pre-warmed to 37°C). The samples were washed four times for 15 min each with 500μL of probe wash buffer (at 37°C) followed by three washes for 5 min each in saline-sodium citrate buffer with Tween. Samples were then incubated in 300μL amplification buffer for 30 min at room temperature. Hairpins h1 and h2 (514nm fluorophore) were mixed at a concentration of 30pmol each, heated at 95 °C for 90 seconds and snap-cooled to room temperature. After 30 mins at room temperature, the hairpins were mixed with 300μL amplification buffer. Samples were incubated overnight in the dark in the hairpin solution. Following this incubation, hairpin solution was removed through six 10-minute 5x SSCT washes followed by two 5-minute PBST washes at room temperature. HCR samples were then counterstained in a 10ng/ml Propidium Iodide (PI, Invitrogen) solution in PBST for 24h at 4°C. The PI solution was then washed in PBST for five minutes at room temperature, followed by an overnight PBST wash at 4°C. Samples were stored in PBST at 4 °C until microscopy.

### Live cell microscopy of embryonic and larval zebrafish

Animals were anaesthetized with 0.2 mg/ml MS-222 (PHARMAQ, UK). For confocal microscopy, animals were mounted laterally in 1.5% ultrapure low melting point agarose (Invitrogen) on a glass coverslip. The coverslip was flipped over on a glass slide with a ring of high-vacuum grease filled with a drop of Danieau’s solution to prevent drying out of the agarose. After imaging, the animals were either sacrificed or released from the agarose using microsurgery blades and kept individually until further use.

For microscopy, 12-bit confocal images were acquired on Leica TCS SP8 laser scanning microscopes. We used 448 nm wavelength for excitation of Cerulean; 488 nm for EGFP and Alexa Fluor(AF) 488; 514 nm for EYFP and AF514; 552nm for mScarlet and KillerRed. For overview images and analysis of cell numbers (i.e. nuclear transgenes), we used 10x/0.4NA (acquisition with 568nm pixel size (xy), 2 µm z-spacing) and 20x/0.7NA (acquisition with 142nm pixel size (xy), 1 µm z-spacing) objectives. Time lapse imaging was performed using a 10x/0.4NA objective (acquisition every 15 mins with 284 pixel size (xy), 5 µm z-spacing). For all other analysis, images were acquired using a 25x/0.95NA H_2_O immersion objective with 114-151nm pixel size (xy) and 1 µm z-spacing. When images were acquired for subsequent deconvolution, x/y/z parameters were increased closer to Nyquist resolution to be compatible for processing with Huygens software (SVI).

### Analysis and presentation of imaging data

All imaging data were analysed using Fiji ^51^, Microsoft Excel and GraphPad Prism. Counting of cells was performed using the Cell Counter plugin for Fiji. Sholl analyses were performed by tracing OPC processes using the SNT plugin on Fiji then running Sholl analysis, which drew concentric circles 2μm apart around the cell soma. 3D rotations were also perfomed on Fiji using the 3D projection tool.

### Statistics and Reproducibility

Zebrafish larvae were derived from the same clutches and pre-selected for appropriate transgene expression where required. Following this initial screen, animals were allocated to treatment groups at random. For the optical treadmill experiments, the response of animals was visually inspected during the first two hours of chronic stimulation, and any non-responsive animals were removed from the experiment. No other data were excluded. Statistical analysis was not used to pre-determine sample sizes. We selected sample sizes based on similar sample sizes that we and others have previously reported for similar experiments ^14,46,52^ . Statistical analyses were performed with Microsoft Excel and GraphPad Prism. All data were tested for normal distribution using the Shapiro-Wilk normality test before statistical testing. For normally distributed data, paired and unpaired t-tests were performed, as appropriate. For data that were not normally distributed, Mann-Whitney U tests were used. The Sholl analysis data was tested using a mixed effects analysis. In the figures, data are shown as mean ± median interquartile range, mean ± standard deviation, mean ± 95% confidence interval, or box and whisker plots as indicated in the figure legends.

## Data availability

All data underlying this study will be made available upon reasonable request. Raw sequence data, gene expression and cell type annotation tables of bulk RNA sequencing data of zebrafish OPCs have been deposited in the Gene Expression Omnibus (GEO) with the accession number GSE317464. The code used for tracking of motor activity is available at: https://doi.org/10.5281/zenodo.18345729

## References

1. Simons, M., Gibson, E. M. & Nave, K.-A. Oligodendrocytes: Myelination, Plasticity, and Axonal Support. Cold Spring Harb. Perspect. Biol. a041359 (2024) doi:10.1101/cshperspect.a041359.

2. Miller, D. J. et al. Prolonged myelination in human neocortical evolution. PNAS 109, 16480–16485 (2012).

3. Hill, R. A., Li, A. M. & Grutzendler, J. Lifelong cortical myelin plasticity and age-related degeneration in the live mammalian brain. Nature Neuroscience 21, 683–695 (2018).

4. Monje, M. Myelin Plasticity and Nervous System Function. Annu Rev Neurosci 41, 61–76 (2018).

5. Xin, W. & Chan, J. R. Myelin plasticity: sculpting circuits in learning and memory. Nat Rev Neurosci 21, 682–694 (2020).

6. Franklin, R. J. M. & ffrench-Constant, C. Regenerating CNS myelin - from mechanisms to experimental medicines. Nature Reviews Neuroscience 18, 753–769 (2017).

7. Dawson, M. R. L., Polito, A., Levine, J. M. & Reynolds, R. NG2-expressing glial progenitor cells: an abundant and widespread population of cycling cells in the adult rat CNS. Mol Cell Neurosci 24, 476–488 (2003).

8. Bergles, D. E. & Richardson, W. D. Oligodendrocyte Development and Plasticity. Csh Perspect Biol 8, a020453 (2015).

9. Hill, R. A., Nishiyama, A. & Hughes, E. G. Features, Fates, and Functions of Oligodendrocyte Precursor Cells. Cold Spring Harb. Perspect. Biol. a041425 (2023) doi:10.1101/cshperspect.a041425.

10. Hughes, E. G. & Stockton, M. E. Premyelinating Oligodendrocytes: Mechanisms Underlying Cell Survival and Integration. Frontiers Cell Dev Biology 9, 714169 (2021).

11. Bhambri, A. et al. Genetic targeting of premyelinating oligodendrocytes reveals activity-dependent myelination mechanisms. Nat. Neurosci. 1–16 (2025) doi:10.1038/s41593-025-02110-1.

12. Viganò, F., Möbius, W., Götz, M. & Dimou, L. Transplantation reveals regional differences in oligodendrocyte differentiation in the adult brain. Nature Neuroscience 16, 1370–1372 (2013).

13. Spitzer, S. O. et al. Oligodendrocyte Progenitor Cells Become Regionally Diverse and Heterogeneous with Age. Neuron 101, 1–13 (2019).

14. Marisca, R. et al. Functionally distinct subgroups of oligodendrocyte precursor cells integrate neural activity and execute myelin formation. Nature Neuroscience 23, 363–374 (2020).

15. Chapman, T. W., Olveda, G. E., Bame, X., Pereira, E. & Hill, R. A. Oligodendrocyte death initiates synchronous remyelination to restore cortical myelin patterns in mice. Nat. Neurosci. 26, 555–569 (2023).

16. Hill, R. A., Patel, K. D., Goncalves, C. M., Grutzendler, J. & Nishiyama, A. Modulation of oligodendrocyte generation during a critical temporal window after NG2 cell division. Nature Neuroscience 17, 1518–1527 (2014).

17. McKenzie, I. A. et al. Motor skill learning requires active central myelination. Science 346, 318–322 (2014).

18. Xiao, L. et al. Rapid production of new oligodendrocytes is required in the earliest stages of motor-skill learning. Nature Neuroscience 19, 1210–1217 (2016).

19. Hughes, E. G., Orthmann-Murphy, J. L., Langseth, A. J. & Bergles, D. E. Myelin remodeling through experience-dependent oligodendrogenesis in the adult somatosensory cortex. Nature Neuroscience 21, 696–706 (2018).

20. Bacmeister, C. M. et al. Motor learning promotes remyelination via new and surviving oligodendrocytes. Nature Neuroscience 23, 819–831 (2020).

21. Miralles, A. J., Unger, N., Kannaiyan, N., Rossner, M. J. & Dimou, L. Analysis of the GPR17 receptor in NG2-glia under physiological conditions unravels a new subset of oligodendrocyte progenitor cells with distinct functions. Glia (2023) doi:10.1002/glia.24356.

22. Orger, M. B., Smear, M. C., Anstis, S. M. & Baier, H. Perception of Fourier and non-Fourier motion by larval zebrafish. Nature Neuroscience 3, 1128–1133 (2000).

23. Chen, S., Chiu, C. N., McArthur, K. L., Fetcho, J. R. & Prober, D. A. TRP channel mediated neuronal activation and ablation in freely behaving zebrafish. Nature Methods 13, 147–150 (2016).

24. Neely, S. A. et al. New oligodendrocytes exhibit more abundant and accurate myelin regeneration than those that survive demyelination. Nat Neurosci 1–6 (2022) doi:10.1038/s41593-021-01009-x.

25. Caterina, M. J. et al. The capsaicin receptor: a heat-activated ion channel in the pain pathway. Nature 389, 816–824 (1997).

26. Gau, P. et al. The zebrafish ortholog of TRPV1 is required for heat-induced locomotion. Journal of Neuroscience 33, 5249–5260 (2013).

27. Czopka, T., ffrench-Constant, C. & Lyons, D. A. Individual oligodendrocytes have only a few hours in which to generate new myelin sheaths in vivo. Developmental Cell 25, 599–609 (2013).

28. Morris, J. K. et al. The 36K protein of zebrafish CNS myelin is a short-chain dehydrogenase. 45, 378–391 (2004).

29. Nagarajan, B. et al. CNS myelin protein 36K regulates oligodendrocyte differentiation through Notch. Glia 68, 509–527 (2020).

30. Marques, S. et al. Oligodendrocyte heterogeneity in the mouse juvenile and adult central nervous system. Science 352, 1326–1329 (2016).

31. Jäkel, S. et al. Altered human oligodendrocyte heterogeneity in multiple sclerosis. Nature 566, 543–547 (2019).

32. Chen, Y. et al. The oligodendrocyte-specific G protein-coupled receptor GPR17 is a cell-intrinsic timer of myelination. Nature Neuroscience 12, 1398–1406 (2009).

33. Wang, J. et al. Robust Myelination of Regenerated Axons Induced by Combined Manipulations of GPR17 and Microglia. Neuron (2020) doi:10.1016/j.neuron.2020.09.016.

34. Häberlein, F. et al. Humanized zebrafish as a tractable tool for in vivo evaluation of pro-myelinating drugs. Cell Chem. Biol. 29, 1541–1555.e7 (2022).

35. Lecca, D., Raffaele, S., Abbracchio, M. P. & Fumagalli, M. Regulation and signaling of the GPR17 receptor in oligodendroglial cells. Glia 68, 1957–1967 (2020).

36. Viganò, F. et al. GPR17 expressing NG2-Glia: Oligodendrocyte progenitors serving as a reserve pool after injury. (2015) doi:10.1002/glia.22929.

37. Provost, E., Rhee, J. & Leach, S. D. Viral 2A peptides allow expression of multiple proteins from a single ORF in transgenic zebrafish embryos. Genesis 45, 625–629 (2007).

38. Paez, P. M. & Lyons, D. A. Calcium Signaling in the Oligodendrocyte Lineage: Regulators and Consequences. Annu Rev Neurosci (2020) doi:10.1146/annurev-neuro-100719-093305.

39. Li, J., Fiore, F., Monk, K. R. & Agarwal, A. Spatiotemporal calcium dynamics orchestrate oligodendrocyte development and myelination. Trends Neurosci. 48, 377–388 (2025).

40. Almeida, R. & Lyons, D. Oligodendrocyte Development in the Absence of Their Target Axons In Vivo. Plos One 11, e0164432 (2016).

41. Fancy, S. P. J., Chan, J. R., Baranzini, S. E. & Rowitch, D. H. Myelin Regeneration: A Recapitulation of Development? Annu Rev Neurosci 34, 110301101035033 (2010).

42. Karttunen, M. J., Czopka, T., Goedhart, M., Early, J. J. & Lyons, D. A. Regeneration of myelin sheaths of normal length and thickness in the zebrafish CNS correlates with growth of axons in caliber. PLoS ONE 12, e0178058 (2017).

43. Vagionitis, S. et al. Clusters of neuronal neurofascin prefigure the position of a subset of nodes of Ranvier along individual central nervous system axons in vivo. Cell Reports 38, 110366 (2022).

44. Auer, F., Vagionitis, S. & Czopka, T. Evidence for Myelin Sheath Remodeling in the CNS Revealed by In Vivo Imaging. Current Biology 28, 549-559.e3 (2018).

45. Almeida, R. G., Czopka, T., ffrench-Constant, C. & Lyons, D. A. Individual axons regulate the myelinating potential of single oligodendrocytes in vivo. Development (Cambridge, England) 138, 4443–4450 (2011).

46. Xiao, Y., Petrucco, L., Hoodless, L. J., Portugues, R. & Czopka, T. Oligodendrocyte precursor cells sculpt the visual system by regulating axonal remodeling. Nat. Neurosci. 25, 280–284 (2022).

47. Kwan, K. M. et al. The Tol2kit: a multisite gateway-based construction kit for Tol2 transposon transgenesis constructs. Dev Dyn 236, 3088–3099 (2007).

48. Zhang, Y., Marshall-Phelps, K. & Almeida, R. G. Fast, precise and cloning-free knock-in of reporter sequences in vivo with high efficiency. Development 150, dev201323 (2023).

49. Štih, V., Petrucco, L., Kist, A. M. & Portugues, R. Stytra: An open-source, integrated system for stimulation, tracking and closed-loop behavioral experiments. PLoS computational biology 15, e1006699 (2019).

50. Bindea, G. et al. ClueGO: a Cytoscape plug-in to decipher functionally grouped gene ontology and pathway annotation networks. Bioinformatics (Oxford, England) 25, 1091–1093 (2009).

51. Schindelin, J., et al. Fiji: an open-source platform for biological-image analysis. Nat. Methods 9, 676–682 (2012).

52. Braaker, P. N. et al. Activity-driven myelin sheath growth is mediated by mGluR5. Nat. Neurosci. 28, 1213–1225 (2025).

